# Cestode larvae excite host neuronal circuits via glutamatergic signaling

**DOI:** 10.1101/859009

**Authors:** Anja de Lange, Hayley Tomes, Joshua S Selfe, Ulrich Fabien Prodjinotho, Matthijs B Verhoog, Siddhartha Mahanty, Katherine Smith, William Horsnell, Chummy Sikasunge, Clarissa Prazeres da Costa, Joseph V Raimondo

## Abstract

Neurocysticercosis (NCC) is caused by infection of the brain by larvae of the parasitic cestode *Taenia solium*. It is the most prevalent parasitic infection of the central nervous system and one of the leading causes of adult-acquired epilepsy worldwide. However, little is known about how cestode larvae affect neurons directly. To address this, we used whole-cell patch-clamp electrophysiology and calcium imaging in rodent and human brain slices to identify direct effects of cestode larval products on neuronal activity. We found that both whole cyst homogenate and excretory/secretory products of cestode larvae have an acute excitatory effect on neurons, which can trigger seizure-like events *in vitro*. Underlying this effect was cestode*-*induced neuronal depolarization, which was mediated by glutamate receptor activation but not by nicotinic acetylcholine receptors, acid-sensing ion channels nor Substance P. Glutamate-sensing fluorescent reporters (iGluSnFR) and amino acid assays revealed that the larval homogenate of the cestodes *Taenia crassiceps* and *Taenia solium* contained high concentrations of the amino acids glutamate and aspartate. Furthermore, we found that larvae of both species consistently produce and release these excitatory amino acids into their immediate environment. Our findings suggest that perturbations in glutamatergic signaling may play a role in seizure generation in NCC.

## Introduction

Neurocysticercosis (NCC) is the most prevalent parasitic infection of the central nervous system (CNS) (1,2). It is caused by the presence of larvae of the cestode *Taenia solium (T. solium)* in the brain (3). The most common symptom of NCC is recurrent seizures (4). As a result, NCC is one of the leading causes of adult-acquired epilepsy worldwide (5), resulting in significant morbidity and mortality. In *T.solium* endemic areas, 29% of people with epilepsy also had NCC (6). Despite the impact of NCC, there are a paucity of studies investigating the seizure mechanisms involved (7). As far as we are aware, to date there have been no studies investigating the direct, acute effects of cestode larval products on neurons. As a result, precisely how larvae perturb neuronal circuits is still relatively poorly understood.

In NCC seizures may occur at any stage following initial infection (8). It is thought that inflammatory processes in the brain play an important role in the development of recurrent seizures (9). Previous research investigating seizure development in NCC has therefore typically focused on how the host neuroinflammatory response to larvae might precipitate seizures (7,10). One study by Robinson *et al*. found that production of the inflammatory molecule and neurotransmitter, Substance P, produced by peritoneal larval granulomas, precipitated acute seizures (11). In addition to host-derived substances, cestodes themselves are known to excrete or secrete various products that interact with host cells in their vicinity. Cestode larvae-derived factors are known to modulate the activation status of immunocytes such as microglia and dendritic cells (12,13). However, comparatively little is known about how factors contained in, or secreted by, cestode larvae might affect neurons and neuronal networks directly, including whether these may have pro-seizure effects.

To address this, we used whole-cell patch-clamp recordings and calcium imaging in rodent hippocampal organotypic slice cultures and in acute human cortical brain slices to demonstrate the direct effects of larval products on neuronal activity. We find that both the whole cyst homogenate and the excretory/secretory (E/S) products of *Taenia crassiceps (T. crassiceps)* and *T. solium* larvae have a strong, acute excitatory effect on neurons. This was sufficient to trigger seizure-like events (SLEs) *in vitro*. Underlying SLE induction was larval induced neuronal depolarization, which was mediated by glutamate receptor activation and not nicotinic acetylcholine receptors, acid-sensing ion channels nor Substance P. Both imaging using glutamate-sensing fluorescent reporters (iGluSnFR) and direct measurements using amino acid assays revealed that the homogenate and E/S products of both *T. crassiceps* and *T. solium* larvae contain high levels of the excitatory amino acid glutamate and to a lesser extent aspartate. Lastly, we provide evidence that larvae of both species, to varying degrees, can produce and release these excitatory amino acids into their immediate environment. This suggests that these parasites release amino acids that could contribute to seizure generation in NCC.

## Results

### Taenia crassiceps homogenate excites neurons and can elicit epileptiform activity

To investigate the potential acute effects of *T. crassiceps* larvae on neurons, *T. crassiceps* larval somatic homogenate was prepared using larvae, harvested from the peritonea of mice, which were freeze-thawed and homogenized (see Materials and Methods and **Fig. 1A**). Whole-cell patch-clamp recordings were made from CA3 pyramidal neurons in rodent hippocampal organotypic brain slice cultures and layer 2/3 pyramidal neurons in human cortical acute brain slices. Pico-litre volumes of *T. crassiceps* homogenate were directly applied to the soma of neurons using a custom-built pressure ejection system (**Fig. 1A**) (14). Application of the homogenate (20 ms puff) elicited immediate, transient depolarization of the membrane voltage in recordings from rat, mouse and human neurons (**Fig. 1B, C, D**). Increasing the amount of homogenate delivered by increasing the pressure applied to the ejection system resulted in increasingly large membrane depolarization, which could trigger single or multiple action potentials (**Fig. 1B, C, D**). To further explore the acute excitatory effect of *T. crassiceps* on neurons, neuronal networks and the propagation of network activity, we performed fluorescence Ca^2+^ imaging in mouse hippocampal organotypic brain slice cultures. Neurons were virally transfected with the genetically encoded Ca^2+^ reporter, GCAMP6s, under the synapsin promoter and imaged using widefield epifluorescence microscopy (**Fig. 1E** and Materials and Methods). To simulate a pro-ictal environment, a low Mg^2+^ aCSF was used (0.5 mM Mg^2+^) and neurons in the dentate gyrus were imaged whilst small, spatially restricted puffs of *T. crassiceps* homogenate were delivered every 15 s using a glass pipette (**Fig. 1E**). The cells within the direct vicinity of the puffing pipette showed a sharp increase in fluorescence immediately following the delivery of *T. crassiceps* homogenate (**Fig. 1F t_1_**) for all 3 puffs, indicating Ca^2+^ entry following membrane depolarization and action potential generation. Cells in the periphery showed notable increases in fluorescence at a delayed interval following some (but not all) puffs (**Fig. 1F t_2_**). The excitation of these cells are likely the result of being synaptically connected to the cells that were exposed to the puff itself. Indeed, a current-clamp recording from a neuron in the same slice (**Fig. 1F, inset**) indicates that a single, spatially restricted puff of *T. crassiceps* homogenate can elicit the onset of a regenerative seizure-like event lasting far longer than the puff itself.

**Figure 1:**
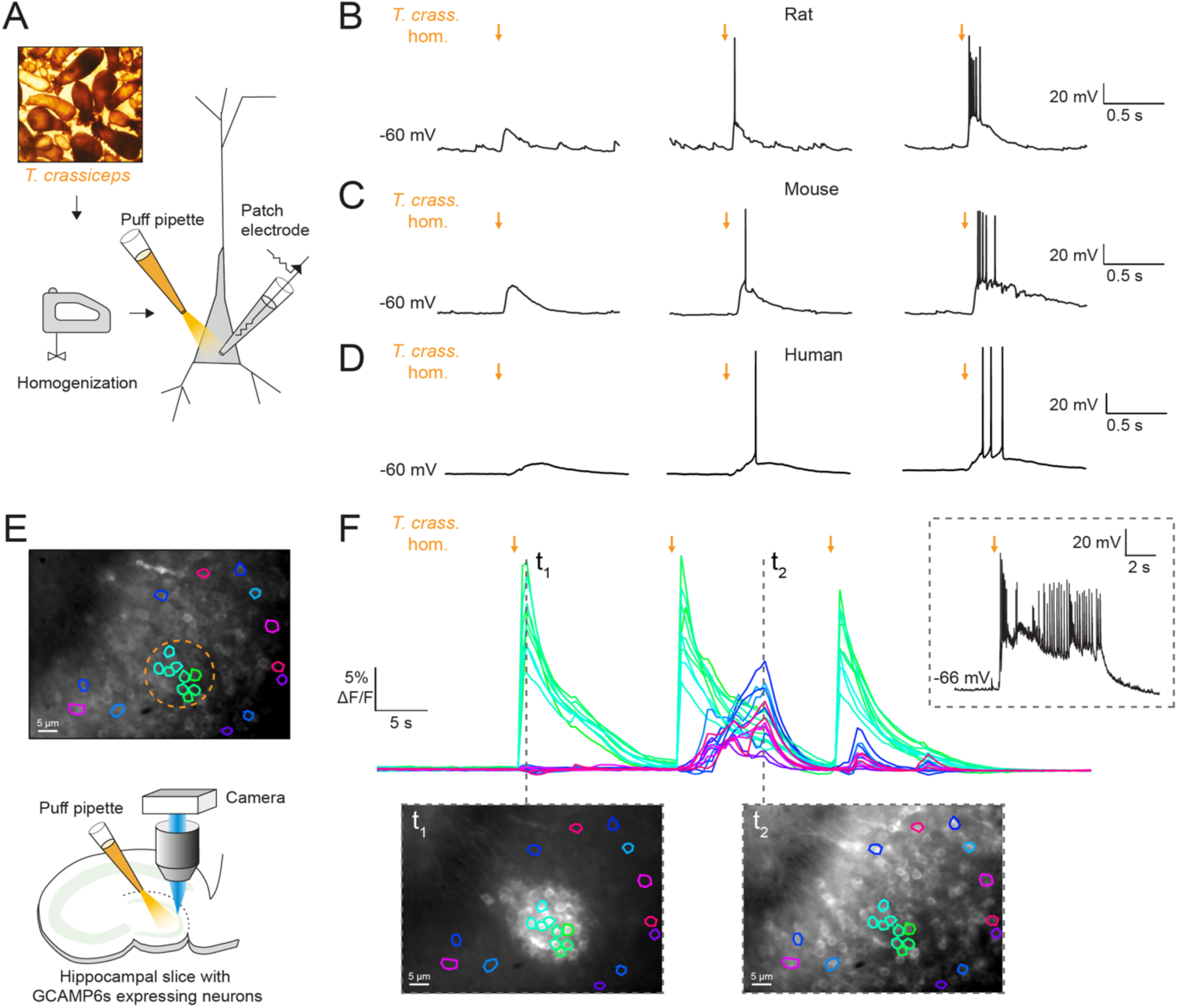
Taenia crassiceps homogenate excites neurons and can elicit epileptiform activity. A) Schematic showing the experimental set up in which whole-cell patch-clamp recordings were made from CA3 hippocampal pyramidal neurons in rodent organotypic slice cultures while a puff pipette delivered pico-litre volumes of homogenised *Taenia crassiceps* larvae (*T. crass.* hom.) targeted at the cell soma. B) Current-clamp recording from a rat pyramidal neuron while increasing amounts of *T. crass.* hom. was applied via the puff pipette (left to right, orange arrows). Small amounts of *T. crass.* hom. resulted in depolarization (left), increasing amounts by increasing the pressure applied to the ejection system elicited single (middle) or even bursts of action potentials (right). C) As in ‘B’, identical effects of *T. crass.* hom. could be elicited in current-clamp recordings from a CA3 hippocampal pyramidal neuron in a mouse organotypic brain slice culture. D) As in ‘B’ and ‘C’, identical effects of *T. crass.* hom. could be elicited in current-clamp recordings from a human frontal lobe layer 2/3 cortical pyramidal neuron in an acute brain slice. E) Top: widefield fluorescence image of neurons in the dentate gyrus of a mouse hippocampal organotypic brain slice culture expressing the genetically encoded Ca^2+^ reporter GCAMP6s under the synapsin promoter in aCSF containing 0.5 mM Mg^2+^. A subset of neurons used to generate the Ca^2+^ traces in ‘E’ are indicated by different colours. The orange dotted circle indicates where *T. crass.* hom. was delivered using the puff pipette. Bottom: schematic of the experimental setup including puff pipette and CCD camera for Ca^2+^ imaging using a 470 nm LED. F) Top, dF/F traces representing Ca^2+^ dynamics from the GCAMP6s expressing neurons labelled in ‘E’ concurrent with 3 puffs (30 ms duration) of *T. crass.* hom. 15 s apart. Bottom: two images of raw Ca^2+^ fluorescence at two time points t_1_ and t_2_. Note how at time point t_2_ neurons distant to the site of *T. crass.* hom. application are also activated, indicating spread of neuronal activity. Inset, top-right: current-clamp recording from a neuron in the region of *T. crass.* hom. application demonstrates seizure-like activity in response to *T. crass.* hom. application.

Most cell types tend to have a high [K^+^]_i_, therefore it is conceivable that the *T. crassiceps* homogenate could have a high K^+^ concentration, which could potentially account for its depolarizing and excitatory effects on neurons. To address this, we directly measured the ionic composition of the *T. crassiceps* homogenate using a Roche Cobas 6000. The K^+^ concentration of the *T. crassiceps* homogenate was 11.4 mM (N = 1, homogenate prepared from >100 larvae) as compared to 3.0 mM in our standard aCSF (N = 1). Direct application of 11.4 mM K^+^ via puff pipette during whole-cell current-clamp from rat CA3 pyramidal neurons resulted in a modest median positive shift of only 0.72 mV (IQR 0.51 - 1.04 mV, N = 8 from 4 slices, **Fig 2B, C**) in the membrane potential, which was lower than the depolarization caused by puffs of *T. crassiceps* homogenate (median 10.12 mV, IQR 9.93 - 12.49 mV, N = 5 from 4 slices), although this did not reach statistical significance (p = 0.26, Kruskal-Wallis test with Dunn’s multiple comparison test, **Fig. 2B, C**). Next, to isolate neuronal depolarization induced by *T. crassiceps* homogenate, but not mediated by K^+^, a cesium based internal solution was used. Despite the addition of cesium the *T. crassiceps* homogenate still resulted in a large depolarisation of the membrane potential (median 13.76, IQR 10.86 - 17.24 mV, N = 49 from 48 slices) which was larger than that caused by 11.4 mM K^+^ (p < 0.0001, Kruskal-Wallis test with Dunns’ multiple comparison test, **Fig. 2B, C**) but was not statistically different from that caused by *T. crassiceps* homogenate without cesium internal solution (p = 0.25, Kruskal-Wallis test with Dunns’ multiple comparison test, **Fig. 2B, C**). Together, this indicates that although K^+^ in the *T. crassiceps* homogenate may contribute to membrane depolarization, much of the effect is mediated by a different component.

**Figure 2:**
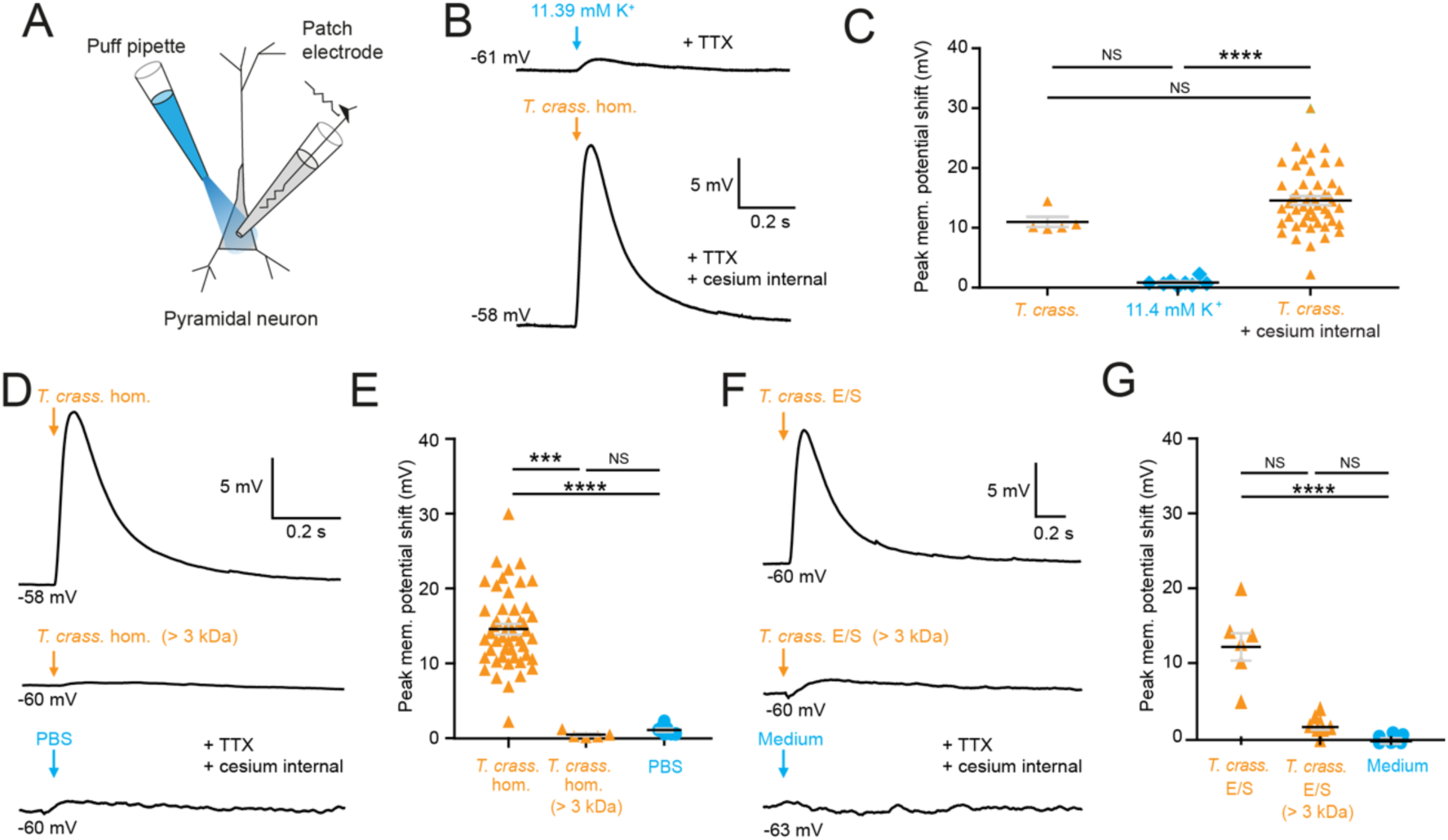
Taenia crassiceps homogenate and E/S product induced neuronal depolarization is mediated by a small molecule. Schematic showing the experimental set up in which whole-cell patch-clamp recordings were made from CA3 hippocampal pyramidal neurons in rat organotypic slice cultures while a puff pipette delivered pico-litre volumes targeted at the cell soma. B) Top trace: 20 ms puff of artificial cerebrospinal fluid (aCSF) containing 11.4 mM K^+^ (equivalent to the K^+^ concentration of *T. crassiceps* homogenate (*T. crass.* hom.)) caused modest depolarization. A standard internal solution was utilised, and 2 μM TTX was added to circulating aCSF to reduce synaptic noise in the voltage trace. Bottom trace: 20 ms puff of *T. crass.* hom. was applied, but a cesium-based internal solution was utilised to block K^+^ channels in the presence of TTX. Puffs of *T. crass* hom. resulted in sizeable depolarization. C) Population data showing that responses to 11.4 mM K^+^ puffs were significantly smaller than that of the *T. crass* hom. when a cesium internal solution was utilised (*T*. *crass*. + cesium internal), but not when a standard internal solution was utilised (*T*. *crass*.). D) Delivery of *T. crass.* hom. caused a depolarizing shift in membrane potential (top trace). The depolarizing response to *T. crass* hom. was largely abolished by dialysing out all molecules smaller than 3 kDa (middle trace). PBS, the solvent for *T. crass.* hom., did not induce a large neuronal depolarisation (bottom trace). E) Population data showing that the membrane depolarization induced by *T. crass.* hom is not due to the PBS solvent and is due to a molecule smaller than 3 kDa. F) Delivery of a puff of *T. crass* excretory/secretory (E/S) products also caused a depolarizing shift in membrane potential (top trace), which was largely abolished by filtering out all molecules smaller than 3 kDa (middle trace). Culture medium, the solvent for the *T. crass* E/S did not induce depolarization (bottom trace). G) Population data showing that the membrane depolarization induced by *T. crass.* E/S is not due to the culture medium solvent and is due to a molecule smaller than 3 kDa. Values with medians ± IQR; *** p ≤ 0.001, ****p ≤ 0.0001, NS = not significant.

To determine the fraction of the *T. crassiceps* homogenate which underlies the acute excitatory effect on neurons we observed, we used a dialysis membrane to separate the fraction of the homogenate bigger than 3 kDa from the total homogenate. This removed molecules smaller than 3 kDa from the homogenate. When this dialysed *T. crassiceps* was puffed onto the cells the depolarizing response was greatly reduced. . The median positive shift in membrane potential for dialysed *T. crassiceps* homogenate was only 0.27 mV (IQR 0.20 - 0.85 mV, N = 5 from 4 slices) as compared to a median for the total homogenate which was 13.76 mV (IQR 10.87 - 17.24 mV, N = 49 from 48 slices, p = 0.0002, Kruskal-Wallis test with Dunn’s multiple comparison test, **Fig. 2D, E**). Phosphate buffered saline (PBS), the solvent for the homogenate also did not produce substantial depolarisation (median 1.00 mV, IQR 0.58 - 1.56 mV, N = 8 from 2 slices,) when compared to the dialysed *T. crassiceps* homogenate (p > 0.10, Kruskal-Wallis test with Dunn’s multiple comparison test, **Fig 2D, E**). This suggests that the excitatory component of the *T. crassiceps* homogenate is a molecule smaller than 3 kDa.

Given that helminths are well known to excrete or secrete (E/S) products into their immediate environment, we next sought to determine whether the <3kDa small molecule/s from *T. crassiceps* homogenate, which was found to induce neuronal depolarisation, was also produced as an E/S product by *T. crassiceps* larvae. The media was collected over a period of 21 days and either used unaltered (*T. crassiceps* total E/S products) or buffer exchanged into a > 3 kDa fraction using an Amicon stirred cell. Brief application of total *T. crassiceps* E/S products using a soma directed puff was sufficient to cause neuronal depolarization (median 12.40 mV, IQR 9.94 - 14.07 mV,, N = 7 from 3 slices) significantly larger than that of the media control (median -0.61 mV, IQR –0.80 - 0.35 mV, N = 7 from 2 slices, p ≤ 0.0001, Kruskal-Wallis test with Dunn’s multiple comparison test, **Fig. 2F, G**). However, the *T. crassiceps* E/S product fraction larger than 3 kDa (median 1.25 mV, IQR 1.02 - 2.21 mV, N = 8 from 1 slice) did not generate significant neuronal depolarization as compared to media control (p = 0.17, Kruskal-Wallis test with Dunn’s multiple comparison test, **Fig. 2F, G**). Together this set of experiments demonstrated that both *T. crassiceps* homogenate and T. *crassiceps* E/S products contain an excitatory component, which is a small molecule.

### The excitatory effects of Taenia crassiceps are mediated by glutamate receptor activation

Robinson *et al* (2012) have found Substance P, an abundant neuropeptide and neurotransmitter, in close vicinity to human NCC granulomas (11). We therefore investigated whether Substance P could elicit a similar neuronal depolarizing response to that of *T. crassiceps* homogenate. However, we found that 100 µM Substance P had no acute effect on the membrane potential of CA3 hippocampal pyramidal neurons (median 0.08 mV, IQR 0.01 – 0.12 mV, N = 5 from 1 slice, p = 0.0625, Wilcoxon signed rank test with theoretical median, **Fig. 3A, B**).

**Figure 3:**
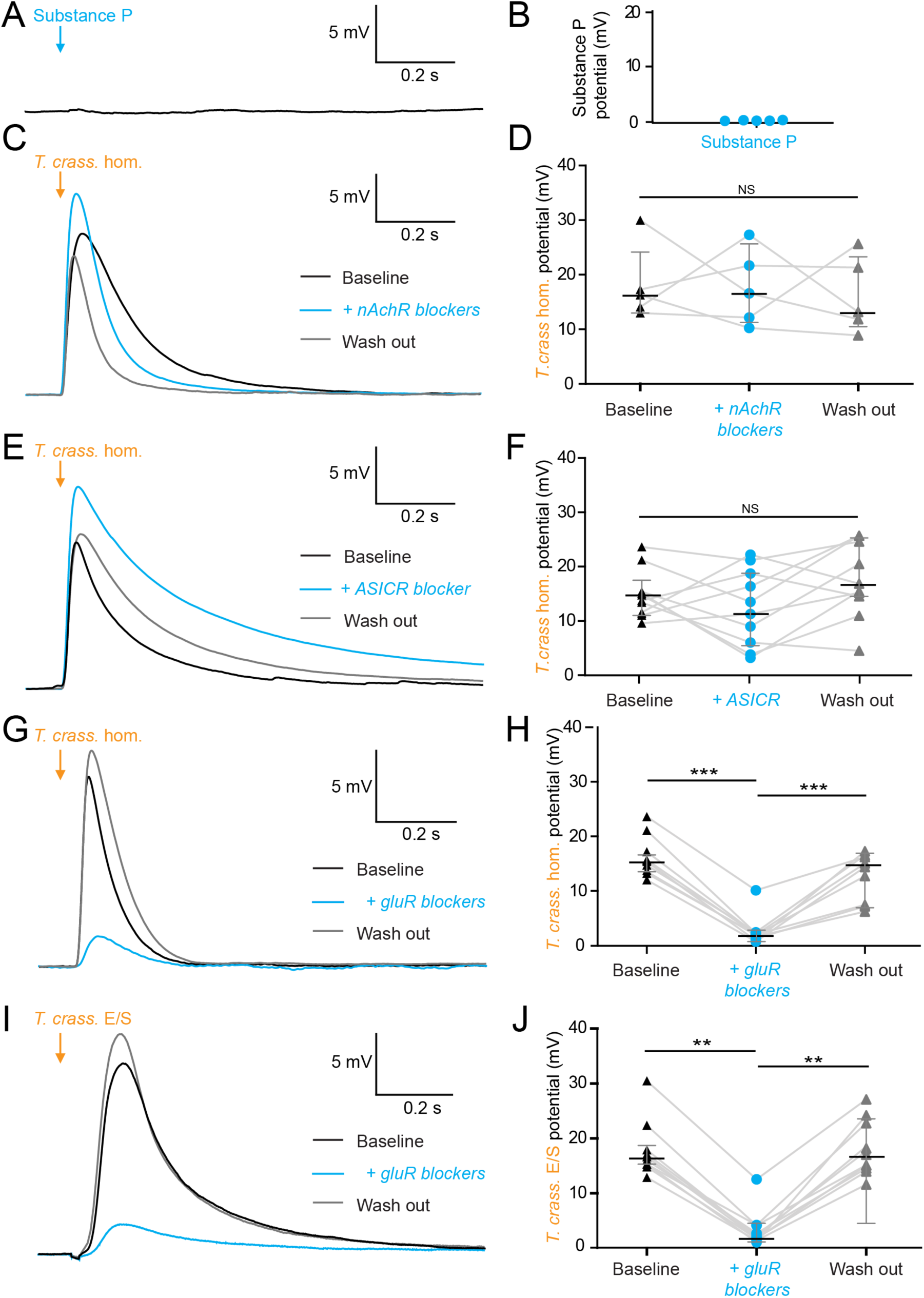
The excitatory effects of Taenia crassiceps are mediated by glutamate receptor activation. Whole-cell patch-clamp recordings in current-clamp were made from CA3 pyramidal neurons in rat organotypic hippocampal slices using a cesium-based internal in the presence of 2 μM TTX. A) 20 ms puff of Substance P (100 µM) via a puff pipette directed at the soma did not affect neuronal membrane potential. B) Population data for Substance P application. C) The depolarization in response to a *T. crassiceps* homogenate (*T. crass.* hom.) puff before (black trace), in the presence of (blue trace), and following wash out (grey trace), of a nicotinic acetylcholine receptor (nAchR) blocker (mecamylamine hydrochloride, 10 µM). D) Population data demonstrating that nAchR blockade has no significant effect on *T. crass.* hom. induced depolarization, NS = not significant. E) The depolarization in response to a *T. crass* hom. puff before (black trace), in the presence of (blue trace), and following washout (grey trace), of an ASIC receptor blocker (amiloride, 2 mM). F) Population data demonstrating that ASIC receptor blockade has no significant effect on *T. crass.* hom. induced depolarization, NS = not significant. G) The depolarization in response to a *T. crass.* hom. puff before (black trace), in the presence of (blue trace), and following washout (grey trace), of a pharmacological cocktail to block glutamate receptors (10µM CNQX, 50µM D-AP5 and 2 mM kynurenic acid). H) Population data showing that the depolarization response to *T. crass.* hom. is significantly reduced in the presence of glutamate receptor blockers and returns upon wash out, ***p ≤ 0.001. I) The depolarization in response to a puff of *T. crass* excretory/secretory (E/S) products is also markedly reduced during glutamate receptor blockade. J) Population data showing that the depolarization response to *T. crass.* hom. is significantly reduced in the presence of glutamate receptor blockers and returns upon wash out, **p ≤ 0.01.

Next we tested whether nicotinic acetylcholine receptors (nAchRs), well described ionotropic receptors in the nervous system (15), could elicit neuronal depolarisation. However, we found that blockade of nAchRs with mecamylamine hydrochloride (10µM) did not significantly alter the *T. crassiceps* homogenate induced depolarisation. The median *T. crassiceps* homogenate induced depolarization was 16.28 mV (IQR 13.54 - 23.63 mV) during baseline, 16.58 mV (IQR 11.21 - 24.50 mV) in the presence of mecamylamine hydrochloride and 13.20 mV (IQR = 10.37 - 23.48 mV) following washout (N = 5 from 5 slices, p = 0.6914, Friedman test, **Fig. 3C, D**). Acid-sensing ion channels (ASICs) are proton-gated sodium channels known to be expressed by hippocampal neurons and result in neuronal depolarization when activated. However, blockade of ASICs with the non-specific ASIC blocker amiloride (2 mM) did not significantly attenuate the effect of the homogenate (**Fig. 3E**), with the median *T. crassiceps* homogenate induced depolarization being 13.86 mV (IQR 11.00 – 16.48 mV) during baseline, 11.99 mV (IQR 4.97 – 19.03 mV) in the presence of amiloride, and 15.79 mV (IQR 13.25 – 24.67 mV) following washout (N = 10 from 10 slices, p = 0.1873, Friedman test, **Fig. 3E, F**).

Ionotropic glutamate receptors (GluRs) including α-amino-3-hydroxy-5-methyl-4-isoxazoleproprionic acid (AMPA), Kainate and N-methyl-D-aspartate (NMDA) receptors (16) are the major source of fast neuronal depolarisation in neurons, therefore it is likely that *T. crassiceps* homogenate could cause neuronal depolarization by activation of GluRs. To test this possibility, *T. crassiceps* homogenate- induced neuronal depolarization was measured before, during and after the application of 10µM CNQX, 50µM D-AP5 and 2mM kynurenic acid (in the aCSF) to block all three classes of GluRs (all in the presence of TTX and using a cesium internal solution). We found that GluR blockade significantly reduced the median *T. crassiceps* homogenate induced neuronal depolarization from 14.72 mV (IQR 13.39 - 15.68 mV) to 1.90 mV (IQR 1.09 - 2.21 mV)), which recovered to a median value of 14.37 mV (IQR 7.32 - 16.38 mV) following washout (N = 9 from 9 slices, p ≤ 0.01, Friedman test with Dunn’s multiple comparison test, **Fig. 3G, H**). Similarly for *T. crassiceps* E/S products we found that application of GluR antagonists reduced the median total *T. crassiceps* E/S products induced depolarization from 16.35 mV (IQR 15.17 - 19.03 mV) to 2.19 mV (IQR 1.67 - 4.16 mV), which recovered to a median value of 16.43 mV (IQR 14.41 - 23.14 mV) following washout (N = 10 from 10 slices, p ≤ 0.01, Friedman test with Dunn’s multiple comparison test, **Fig. 3I, J**). These results indicate that the acute excitatory effects of both *T. crassiceps* homogenate and *T. crassiceps* E/S products are mediated through GluRs.

### Taenia crassiceps products contain excitatory amino acids detected using iGluSnFR

Glutamate is the prototypical agonist of GluRs (16). We therefore sought to directly detect glutamate in *T. crassiceps* larval products using glutamate-sensing fluorescent reporters. We utilized the genetically encoded glutamate reporter (iGluSnFR), which was virally transfected into mouse organotypic hippocampal slice cultures under the synapsin 1 gene promoter and imaged using widefield epifluorescence microscopy (**Fig. 4A** and Materials and Methods). Brief puffs of *T. crassiceps* homogenate delivered over the soma of iGluSnFR-expressing pyramidal neurons caused a robust increase in fluorescence (median 23.32 %, IQR 5.79 – 37.66 %, N = 35 from 6 slices, **Fig. 4B, G**) when compared to puffs of aCSF (median 0.0012 %, IQR -0.42 – 1.02 %, N = 12 from 1 slice, p < 0.0001, Kruskal-Wallis test with Dunn’s multiple comparison test, **Fig. 4G**). Fluorescence increases were also observed with puffs of *T. crassiceps* E/S products (median 11.69 %, IQR 2.94 – 15.85 %, N = 10 from 2 slices, **Fig. 4C, G**) although this did not reach statistical significance when compared to puffs of aCSF (median 0.0012 %, IQR -0.42 – 1.02 %, N = 12 from 1 slice, p = 0.16, Kruskal-Wallis test with Dunn’s multiple comparison test, **Fig. 4G**). Aspartate has a similar chemical structure to glutamate and is also known to excite neurons via glutamate receptor activation (17). iGluSnFR is also known to be sensitive to aspartate as well as glutamate (18). We found that puffing aCSF containing 100 μM glutamate (median 31.35 %, IQR 15.35 – 54.92 %, N = 14 from 2 slices, **Fig. 4E, G**) or 100 μM aspartate (median 15.09 %, IQR 11.12 – 33.12 %, N = 5 from 1 slice, **Fig. 4F, G**) onto hippocampal pyramidal neurons expressing iGluSnFR resulted in a significantly higher fluorescence as compared to aCSF (median 0.0012 %, IQR -0.42 – 1.02 %, N = 12 from 1 slice, Kruskal-Wallis test with Dunn’s multiple comparison test, p ≤ 0.05, **Fig. 4G**). Similarly, puffing aCSF containing 100 μM glutamate (median 15.05 mV, IQR 11.12 – 18.71 mV, N = 6 from 1 slice, **Fig. 4H, I**) or 80 μM aspartate (median 7.75 mV, IQR 3.74 – 15.99 mV, N = 5 from 1 slice, **Fig. 4H, I**) during whole-cell current-clamp recordings elicited membrane depolarizations which were similar those observed with *T. crassiceps* homogenate (median 13.76 mV, IQR 10.87 – 17.24 mV, N = 49 from 48 slices, **Fig. 4I**) and E/S products (median 12.40 mV, IQR 9.94 – 14.07 mV, N = 7 from 3 slices, Kruskal-Wallis test with Dunn’s multiple comparison test, p > 0.05, **Fig. 4I**).

**Figure 4:**
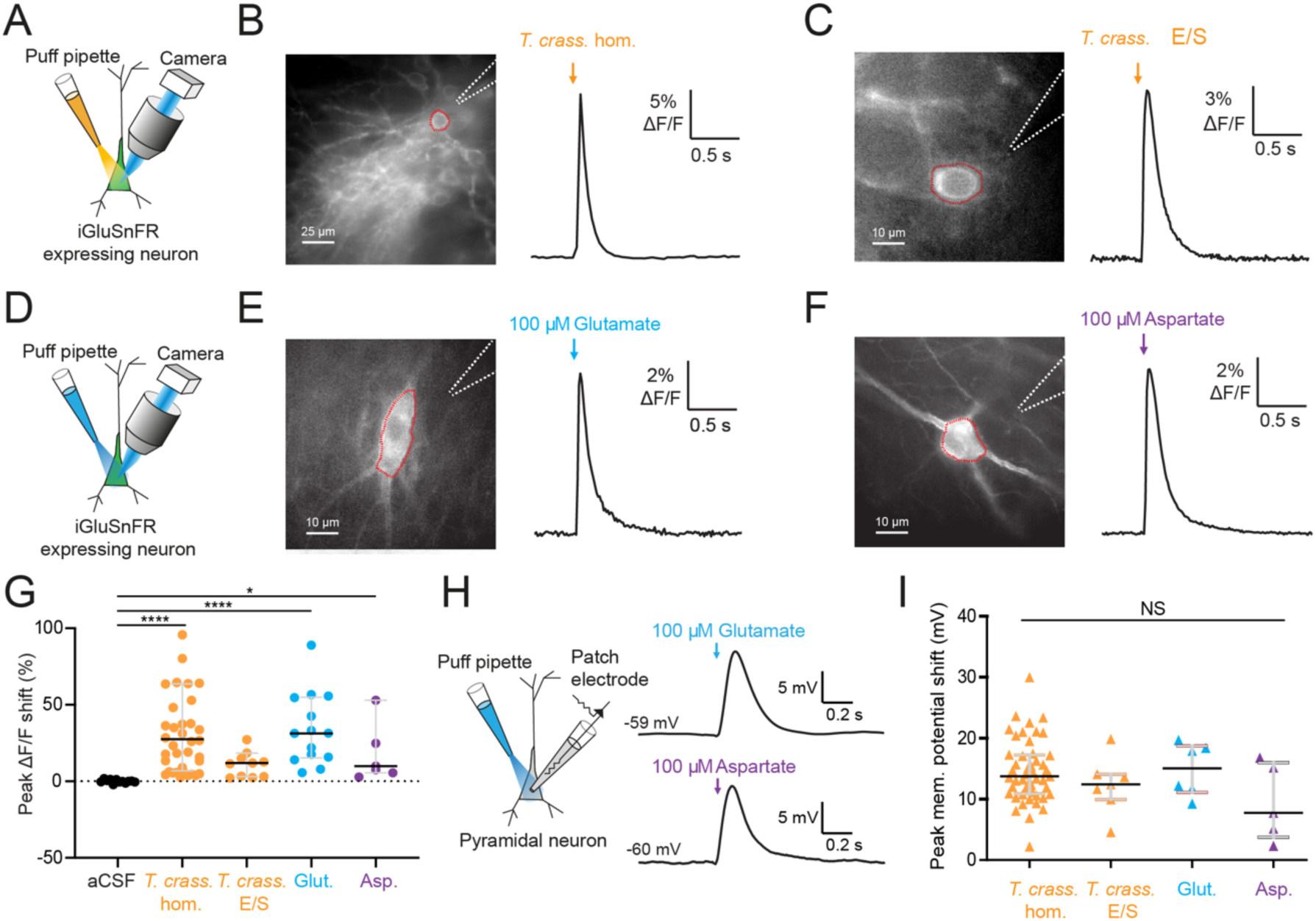
Taenia crassiceps products contain excitatory amino acids detected using iGluSnFR. A) Schematic of the experimental setup for glutamate detection by the genetically encoded fluorescent glutamate reporter iGluSnFR, including puff pipette and sCMOS camera for imaging following excitation using a 475/28 nm LED-based light engine. B) Left, iGluSnFR expressing neuron with the region of interest (red dashed line) used to calculate dF/F trace (right) during a 20 ms *T. crassiceps* homogenate (*T. crass.* hom.) puff (orange arrow). C) As in ‘B’ but with a 20 ms *T. crassiceps* excretory/secretory (*T. crass.* E/S) product puff (orange arrow). D) Schematic of experimental setup as in ‘A’ but with glutamate or aspartate application via puff pipette. E) iGluSnFR fluorescence just after a 20 ms puff of aCSF containing 100 μM glutamate (blue arrow). F) As in ‘E’ but with iGluSnFR fluorescence during a 20 ms puff of aCSF containing 100 μM aspartate (purple arrow). G) Population data comparing peak dF/F shifts after a 20 ms puff of aCSF (as a negative control), *T. crass.* hom., *T. crass.* E/S products, aCSF containing 100 μM glutamate (as a positive control), or aCSF containing 100 μM aspartate (as a positive control), values with medians ± IQR, * p ≤ 0.05, **** p ≤ 0.0001. H) Schematic of whole-cell patch-clamp recording in current-clamp mode from a CA3 pyramidal neuron using a cesium-based internal and in the presence of 2 μM TTX. 20 ms puff of aCSF containing 100 μM glutamate produces a significant depolarizing shift in membrane potential (top trace), a similar puff but with 100 μM aspartate elicited a similar response (bottom trace). I) Population data comparing peak membrane potential shift after a 20 ms puff of *T. crass.* hom., *T. crass.* E/S, aCSF containing 100 μM glutamate, and aCSF containing 100 μM aspartate. Values with medians ± IQR; NS = not significant.

### Taenia crassiceps larvae contain and produce glutamate and aspartate

Given the short-term responses to homogenate and E/S products we observed using iGluSnFR, we next sought to record possible evidence of changes in ambient glutamate or aspartate using a more naturalistic setting where iGluSnFR-expressing neurons were placed adjacent to a live *T. crassiceps* larva (see Materials and Methods, **Fig. 5A-C**). During imaging periods lasting up to 15 minutes, fluorescence was stable, indicating no detectable oscillatory changes in ambient glutamate/aspartate on the timescales recorded (N = 2 slices, **Fig. 5B, C**). Our submerged recording setup might have led to swift diffusion or washout of released glutamate, possibly explaining the lack of observable changes. We then directly measured the concentration of glutamate and aspartate in our *T. crassiceps* products using separate glutamate and aspartate assays (see Materials and Methods). We found that the median concentration of glutamate in the *T. crassiceps* homogenate and E/S products were 119.9 μM (IQR 73.41 – 153.2 μM, N = 5, **Fig. 5D**) and 72.32 μM (IQR 32.95 – 116.0 μM, N = 6, p = 0.1061, Mann-Whitney test, **Fig. 5D**), respectively. The median concentration of aspartate in *T. crassiceps* homogenate and E/S products were 25.8 μM (IQR 16.5 – 35.1 μM, N = 3, **Fig. 5E**) and 44.03 μM (IQR 17.56 – 84.74 μM, N = 3, p = 0.4000, Mann-Whitney test, **Fig. 5E**), respectively. Together, these findings suggest that *T. crassiceps* larval homogenate and E/S products contain the excitatory amino acids glutamate and aspartate at concentrations that are sufficient to elicit neuronal depolarization.

**Figure 5:**
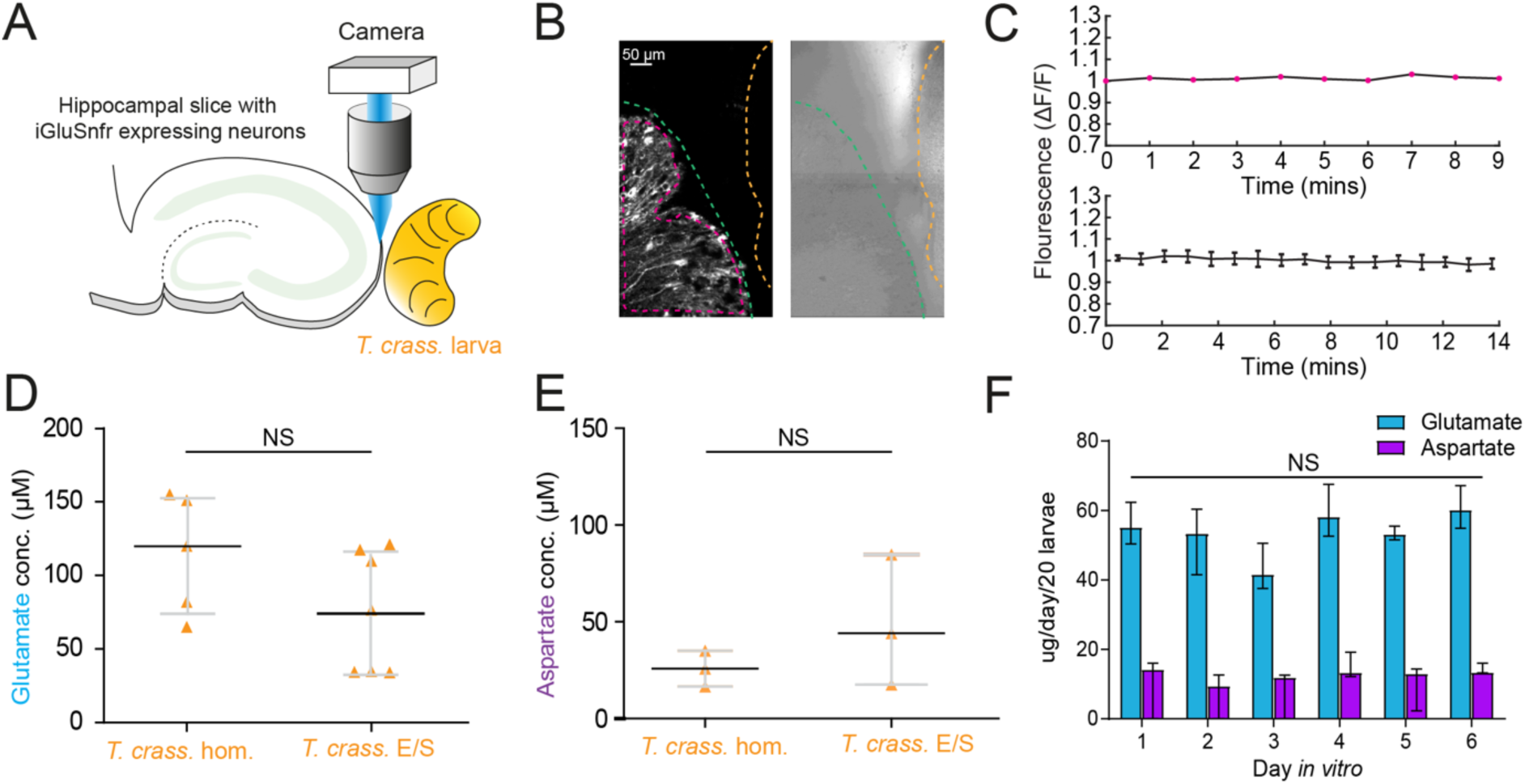
Taenia crassiceps larvae contain and produce glutamate and aspartate. A) Schematic of the experimental setup whereby a living *Taenia crassiceps* (*T. crass.)* larva is placed in close proximity to a hippocampal organotypic brain slice with iGluSnFR-expressing neurons, for detection of glutamate fluctuations. B) Left, fluorescence image of a slice with iGluSnFR-expressing neurons (green dashed line) adjacent to a *T. crass.* larva (orange dashed line), also visible in the transmitted light image (right). The region of interest (pink dashed line) was used to calculate the top dF/F trace in ‘C’. C) Top, dF/F trace from the example in ‘B’ during a 9 minute recording. Lower trace: population data of the **mean ± SEM** from 5 slice-larva pairs showed no detectable oscillations or changes in glutamate. D) Glutamate concentration in *T. crass.* Homogenate (*T. crass.* hom.) and *T. crass.* excretory/secretory (*T. crass*. E/S) products as measured using a glutamate assay kit, values with medians ± IQR; ns = not significant. E) Aspartate concentration in *T. crass.* hom. and *T. crass.* E/S as measured using an aspartate assay kit, values with medians ± IQR; ns = not significant. F) Population data showing *de novo* production of glutamate and aspartate by by by *T. crass* larvae for six days *in vitro*, values with medians ± IQR; NS = not significant.

Next, to determine whether *T. crassiceps* larvae actively produce and excrete/secrete glutamate and/or aspartate into their environments we measured the *de novo* daily production of glutamate and aspartate by larvae following harvest of live larvae. *T. crassiceps* larvae released a relatively constant median daily amount of glutamate, ranging from 41.59 – 60.15 ug/20 larvae, which showed no statistically significant difference across days one to six (N = 3 per day, p = 0.18, Kruskal-Wallis test, **Fig. 5F**). Similarly, *T. crassiceps* larvae released a relatively constant median daily amount of aspartate, ranging from 9.43 – 14.18 ug/20 larvae, which showed no statistically significant difference across days one to six (N = 3 per day, p = 0.28, Kruskal-Wallis test, **Fig. 5F**).

### Taenia solium larvae depolarize human neurons via the production of glutamate

*T. crassiceps*, whilst a closely related species to *T. solium*, is not the causative pathogen in humans. We therefore sought to determine whether homogenate and E/S products from *T. solium* (the natural pathogen in humans), also have an excitatory effect on human neurons within human brain tissue (the natural host). During whole-cell patch-clamp recordings made from human layer 2/3 pyramidal cells in acute cortical brain slices, pico-litre volumes of *T. solium* larval homogenate and *T. solium* larval E/S products were delivered by a glass pipette. Similarly to *T. crassiceps*, both *T. solium* homogenate (**Fig. 6A**) and *T. solium* E/S product (**Fig. 6A**) caused membrane depolarization, which was sufficient to result in action potential firing.

**Figure 6:**
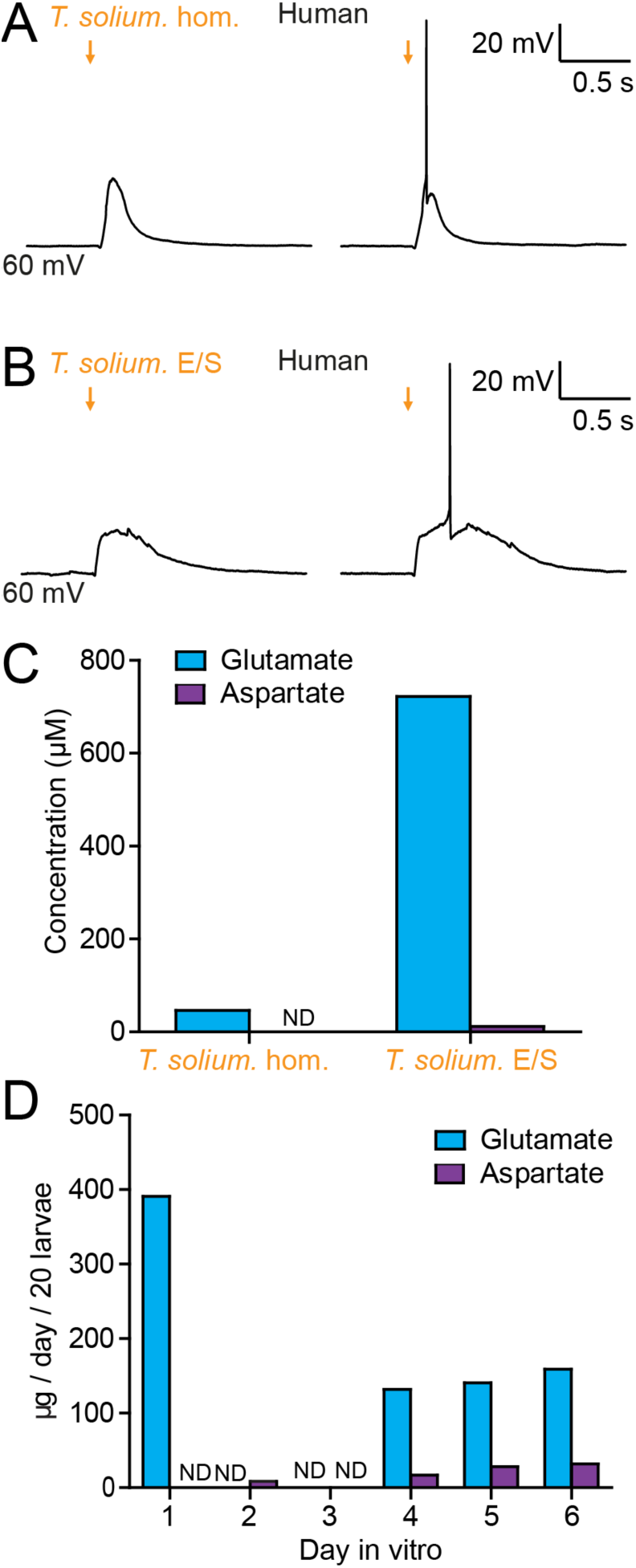
Taenia solium larvae depolarize human neurons via the production of glutamate. A) Whole-cell patch-clamp recordings in current-clamp from a layer 2/3 human frontal lobe cortical pyramidal neuron in an acute brain slice, while increasing amounts of *T. solium* homogenate (*T. solium* hom.) was applied via a puff pipette (left to right, orange arrows). Small amounts of *T. solium* hom. resulted in depolarization (left), while an increased amount elicited an action potential (right). B) As in ‘A’, *T. solium* excretory/secretory (*T. solium* E/S) products elicited membrane depolarization and an increased amount elicited an action potential. C) Concentrations of glutamate and aspartate in *T. solium* hom. and *T. solium* E/S (N = 1, ND = not detectable). D) *De novo* production of glutamate and aspartate by *T. solium* larvae over a six day culture period (N = 1, ND = not detectable).

We then measured the glutamate and aspartate concentration of the *T. solium* homogenate and *T. solium* E/S products. Significant levels of glutamate were detected in both *T. solium* homogenate (46.4 μM, N = 1, **Fig. 6C**) and the *T. solium* E/S product (722 μM, N = 1, **Fig. 6C**). However, aspartate was undetectable in the *T. solium* homogenate (N = 1, **Fig. 6C**) with a low concentration of aspartate observed in the E/S product (15.3 μM, N = 1, **Fig. 6C**). Finally, *T. solium* larvae were observed to release glutamate on the first day post-harvest (391.1 μM/20 larvae, N = 1, **Fig. 6D**) none on the 2^nd^ and 3^rd^ day *in vitro* (N = 1 each, **Fig. 6D)**, but produced significant amounts of glutamate on days 4, 5 and 6 (day 4: 131.9 μM/20 larvae, N = 1; day 5: 140.6 μM/20 larvae, N = 1; day 6: 158.8 μM/20 larvae, N = 1, **Fig. 6D**). Aspartate was not detected on days 1 and 3 (N = 1 each, **Fig. 6D**), but was observed to be released on days 2, 4, 5 and 6 (day 2: 8.4 μM/20 larvae, N = 1; day 4: 16.8 μM/20 larvae, N = 1; day 5: 28.1 μM/20 larvae N = 1; day 6: 31.9 μM/20 larvae, N = 1, **Fig. 6D**). These results demonstrate that *T. solium* larvae continually release glutamate and aspartate into their immediate surroundings.

## Discussion

Here we used patch-clamp electrophysiology, and calcium imaging in rodent hippocampal organotypic slice cultures and human acute cortical slices to demonstrate that cestode larval products cause neuronal depolarization and can initiate seizure-like events via glutamate receptor activation. Glutamate-sensing fluorescent reporters and amino acid assays revealed that *T. crassiceps* and *T. solium* larvae contain and release the excitatory amino acid glutamate and to a lesser extent aspartate.

Clinical evidence and animal models conclusively demonstrate that the presence of cestode larvae in the brain can result in the generation of seizures (10,19). Previous work has focused on the involvement of the host inflammatory response in seizure generation (11,20). Whilst this is likely important, it does not preclude the involvement of additional or exacerbating pathogenic mechanisms for seizure generation and epileptogenesis in NCC. In this study we have identified that *Taenia* larvae have a direct excitatory effect on neurons via glutamatergic signaling. This is important, as the central role of glutamatergic signaling in epileptogenesis has conclusively been shown using multiple cell culture (21–23), slice (24–26), and *in vivo* models of epilepsy (27–29).

Clinically, in addition to NCC, the other major causes of adult acquired epilepsy are stroke, traumatic brain injury and CNS tumors (30,31). In these other causes of acquired epilepsy, glutamatergic signaling and glutamate excitotoxicity are thought to be central to the pathogenic process. Glutamate excitotoxicity occurs when depolarized, damaged or dying neurons release glutamate, which activate surrounding neurons via NMDA and AMPA receptors, resulting in sustained neuronal depolarization, Ca^2+^ influx and the subsequent activation of signaling cascades and enzymes. These in turn lead to cell death via necrosis and apoptosis (32), the further release of intracellular glutamate and the propagation of the excitotoxic process to neighbouring cells. In the neurons that survive, the prolonged exposure to glutamate has been shown to cause hyperexcitability and seizures (21,23) via multiple mechanisms including enhanced intrinsic excitability and NMDA receptor dependent disruption of GABAergic inhibition (33–35). Given our finding that cestode larvae contain and release significant quantities of glutamate, it is possible that homeostatic mechanisms for taking up and metabolizing glutamate fail to compensate for larval-derived glutamate in the extracellular space. Therefore, similar glutamate-dependent excitotoxic and epileptogenic processes that occur in stroke, traumatic brain injury and CNS tumors are likely to also occur in NCC.

Glioma, a common adult primary brain tumor, which typically presents with seizures in over 80% of patients (36), is an intriguing condition for comparison to NCC. Here, the tumor cells themselves have been shown to release glutamate into the extracellular space via the system x_c_^−^cystine-glutamate antiporter (35). Interestingly, there is compelling evidence that tumoral release of glutamate via this mechanism both causes seizures and favours glioma preservation, progression and invasion in cases of malignant glioma (37–39). Analogously it is possible that *Taenia* larvae in the brain utilize the release of glutamate and the induction of glutamate excitotoxicity to facilitate their growth and expansion, with the accompanying effect of seizure generation. In addition, the death of *Taenia* larvae, or larval-derived cells, would also result in the release of metabolic glutamate, further contributing to glutamate release and excitotoxicity. In the case of glioma, where the mechanism of glutamate release by tumor cells is known, pharmacological agents (*e.g.,* sulfasalazine), which block glutamate release have considerable potential as therapeutic agents for reducing seizure burden in this condition. Therefore, it is important that future work attempts to identify the molecular mechanisms underlying the *Taenia*-specific production of glutamate to inform the development of new therapeutic strategies to potentially reduce larval expansion and possibly treat seizures in NCC.

An important consideration is how we might reconcile our findings of glutamate release by *Taenia* larvae with the clinical picture of delayed symptom onset in NCC. In people with NCC, some experience acute seizures immediately following infection whilst others present with seizures months to years following initial infection (40,41). This suggests that multiple different, or possibly interacting, mechanisms might be involved in the epileptogenic process in NCC. A recent report has shown that *T. solium* cyst products mimicking viable cysts contain the enzyme glutamate dehydrogenase, which metabolizes glutamate, whilst products mimicking degenerating cysts do not (42). This predicts that the glutamate levels the brain would be exposed to would be highest when a cyst dies and releases its contents into its immediate surroundings. Neuroinflammation has long been thought to be central to the development of seizures in NCC. In support of this, larval suppression of host inflammatory responses could help explain why many patients may remain asymptomatic for years following infection. Our findings are not at odds with this line of thinking when considering the effect of neuroinflammation on extracellular glutamate uptake. In the healthy brain parenchymal astrocytes are optimized to maintain extracellular glutamate at low concentrations (43). However, the inflammatory transition of astrocytes to a reactive phenotype is known to impede their ability to buffer uptake of extracellular glutamate (44). It is therefore conceivable that cyst-associated reactive astrocytosis may gradually erode the ability of homeostatic mechanisms to compensate for larvae derived extracellular glutamate resulting in delayed symptom onset in NCC.

There is still considerable uncertainty regarding the precise sequence of events leading to seizure onset in NCC. Nonetheless our findings provide the first evidence that, as is the case with the other common causes of adult-acquired epilepsy (*i.e.* stoke, traumatic brain injury and glioma), increased extracellular, parasite-derived excitatory amino acids such as glutamate, and perturbed glutamatergic signaling, possibly also play a role in the development of seizures in NCC.

## Materials and Methods

### Ethics statement

All animal handling, care and procedures were carried out in accordance with South African national guidelines (South African National Standard: The care and use of animals for scientific purposes, 2008) and with authorisation from the University of Cape Town Animal Ethics Committee (Protocol No: AEC 015/015, AEC 014/035). Ethical approval for use of human tissue was granted by the University of Cape Town Human Research Ethics Committee (HREC 016/2018) according to institutional guidelines.

### Taenia maintenance and preparation of whole cyst homogenates and E/S products

Larvae of *T. crassiceps* (ORF strain) were donated to us by Dr Siddhartha Mahanty (University of Melbourne, Melbourne, Australia) and propagated *in vivo* by serial intraperitoneal infection of 5–8- week-old female C57BL/6 mice. Every 3 months parasites were harvested by peritoneal lavage and washed once in phosphate buffered saline (PBS, 1X, pH 7.4) containing penicillin (500 U/ml); streptomycin (500 ug/ml); and gentamicin sulphate (1000 ug/ml) (Sigma-Aldrich), before being washed a further five times in standard PBS (1X).

For the preparation of *T. crassiceps* whole cyst homogenate, larvae were stored immediately after harvesting at -80 °C. Later the larvae were thawed (thereby lysing the cells) and suspended in PBS (1X) containing a protease cocktail inhibitor (1% vol/vol, Sigma-Aldrich) at a larval:PBS ratio of 1:3. The larvae were then homogenised on ice using a Polytron homogenizer (Kinematica). The resulting mixture was centrifuged at 4000 rpm for 20 minutes at 4 °C. The liquid supernatant (between the white floating layer and solid pellet) was collected and sterile filtered through a 0.22 µm size filter (Millex-GV syringe filter, Merck). This supernatant was then aliquoted and stored at -80 °C until use. To assess whether large or small molecules were responsible for the excitation of neurons, a portion of the whole cyst homogenate was dialysed using a Slide-A-Lyzer™ dialysis cassette (3kDa MWCO, Separations) in 2l of artificial cerebro-spinal fluid (aCSF) at 4 °C. The aCSF solution was changed twice over 24 hours. To determine the ionic composition of *T. crassiceps* whole cyst homogenate a Cobas 6000 analyser (Roche) was used, with ion specific electrodes for K^+^ and Na^+^. A Mettler Toledo SevenCompact™ pH meter S210 (Merck) was used to determine the pH of the homogenate.

For the preparation of *T. crassiceps* excretory/secretory products, after harvesting, washed larvae were maintained for 14-21 days at 37 °C in 5 % CO_2_ in 50 ml tissue culture flasks (approximately 7 ml volume of larvae per flask) containing culture medium consisting of Earle’s Balanced Salt Solution (EBSS) supplemented with glucose (5.3 g/liter), Glutamax (1X), penicillin (50 U/ml), streptomycin (50 ug/ml), gentamicin sulphate (100 ug/ml) and nystatin (11.4 U/ml) (Sigma-Aldrich). Culture medium was replaced after 24 hours, and the original media discarded. Thereafter, culture media were replaced and collected every 48-72 hours, stored at -20 °C and at the end of the culture period, thawed and pooled. The pooled conditioned medium was termed total excretory/secretory products (total E/S). A portion of the total E/S was aliquoted and stored at -80 °C, while another portion was concentrated (about 100X) and buffer exchanged to PBS using an Amicon stirred cell with a 3 kDa MWCO cellulose membrane (Sigma-Aldrich). The concentrated product contained *T. crassiceps* E/S products larger in size than 3kDa (E/S > 3kDa), in PBS (1X). This was aliquoted and stored at -80 °C. The fraction of the total E/S that passed through the 3kDa membrane (E/S < 3 KDa, still in medium) was also retained, aliquoted and stored at -80 °C.

For the preparation of *T. solium* whole cyst homogenate, larvae of *T. solium* were harvested from the muscles of a heavily infected, freshly slaughtered pig. After extensive washing with sterile 1X PBS, *T. solium* larvae were suspended in PBS containing phenylmethyl-sulphonyl fluoride (5mM) and leupeptin (2.5 μM) at a larvae:PBS ratio of 1:3. Larvae were then homogenised using a sterile handheld homogenizer at 4 °C. The resulting homogenate was sonicated (4 x 60 s at 20 kHz, 1 mA, with 30 s intervals) and gently stirred with a magnetic stirrer for 2 h at 4 °C. Thereafter it was centrifuged at 15,000 g for 60 min at 4 °C and the liquid supernatant (between the white floating layer and solid pellet) was collected. The supernatant was filtered through 0.45 μm size filters (Millex-GV syringe filter, Merck), aliquoted and stored at -80 °C. All *T. crassiceps* and *T. solium* larval products were assessed for protein concentration using a BCA protein or Bradford’s assay kit respectively.

For the assessment of daily glutamate and aspartate production both *T. crassiceps* and *T. solium* larvae were placed into 6 well plates (+/- 20 per well, of roughly 5mm length) with 2 ml of culture medium (*Taenia solium* medium: RPMI 1640 with 10 mM HEPES buffer, 100 U/ml penicillin, 100 μg/ml streptomycin, 0.25 μg/ml amphotericin B and 2 mM L-glutamine; *Taenia crassiceps* medium: EBSS with 100 U/ml penicillin, 100 μg/ml streptomycin, 11.4 U/ml nystatin and 2 mM Glutamax). Every 24 hours 1 ml of culture medium was collected from each well, stored at -80 °C, and replaced with fresh culture medium. The concentrations of glutamate and aspartate were measured using a Glutamate assay kit and an Aspartate assay kit, respectively, according to the supplier’s instructions (Sigma-Aldrich). ‘*T. solium* E/S’ used in human organotypic brain slice electrophysiology experiments, consisted of a mixture of equal volumes of the *T. solium* medium collected on D1, D2 and D3.

### Brain slice preparation

Rodent hippocampal organotypic brain slices were prepared using 6-8 day old Wistar rats and C57BL/6 mice following the protocol originally described by Stoppini et al., (1991). Brains were extracted and swiftly placed in cooled (4 °C) dissection media consisting of Earle’s balanced salt solution (Sigma-Aldrich); 6.1 g/L HEPES (Sigma-Aldrich); 6.6 g/L D-glucose (Sigma-Aldrich); and 5 µM sodium hydroxide (Sigma-Aldrich). The hemispheres were separated, and individual hippocampi were removed and immediately cut into 350 μm slices using a Mcllwain tissue chopper (Mickle). Cold dissection media was used to rinse the slices before placing them onto Millicell-CM membranes (Sigma-Aldrich). Slices were maintained in culture medium consisting of 25 % (vol/vol) Earle’s balanced salt solution (Sigma-Aldrich); 49 % (vol/vol) minimum essential medium (Sigma-Aldrich); 25 % (vol/vol) heat-inactivated horse serum (Sigma-Aldrich); 1 % (vol/vol) B27 (Invitrogen, Life Technologies) and 6.2 g/l D-glucose. Slices were incubated in a 5 % carbon dioxide (CO_2_) humidified incubator at between 35 – 37 °C. Recordings were made after 6-14 days in culture.

For human cortical acute brain slices, brain tissue from temporal cortex was obtained from patients undergoing elective neurosurgical procedures at Mediclinic Constantiaberg Hospital to treat refractory epilepsy (2 patients, details below).

| <u>Age at surgery</u> | <u>Gender</u> | <u>Reason for surgery</u> | <u>Region</u> | <u>Tissue type</u> | <u>Epilepsy Duration</u> |
| --- | --- | --- | --- | --- | --- |
| 41 | Female | Refractory epilepsy | Temporal Cortex | Access | 2 years |
| 4 | Male | Refractory epilepsy, Malformation of | Temporal Cortex | Least abnormal | unknown |
|  |  | Cortical<br>Development |  |  |  |

Informed consent to use tissue for research purposes was obtained from patients or guardians prior to surgery. Resected brain tissue was transported from surgery to the laboratory in an ice-cold choline-based cutting solution composed of Choline Chloride (110 mM, Sigma-Aldrich); NaHCO_3_ (26 mM, Sigma-Aldrich); D-glucose (10 mM, Sigma-Aldrich); Sodium Ascorbate (11.6 mM, Sigma-Aldrich); MgCl_2_ (7 mM, Sigma-Aldrich); Sodium Pyruvate (3.1 mM, Sigma-Aldrich); KCl (2.5 mM, Sigma-Aldrich); NaH_2_PO_4_ (1.25 mM, Sigma-Aldrich); and CaCl_2_ (0.5 mM, Sigma-Aldrich). Cutting solution was bubbled with carbogen gas (95% O_2_, 5% CO_2_) and had an osmolality of approximately 300 mOsm. In the lab, the pia mater was carefully removed, and tissue was trimmed into blocks for sectioning using a Compresstome (Model VF-210-0Z, Precisionary Instruments). Slices were cut at 350 µm thickness and allowed to recover for ∼30 min at 34 °C in standard aCSF composed of NaCl (120 mM, Sigma-Aldrich); KCl (3 mM, Sigma-Aldrich); MgCl_2_ (2 mM, Sigma-Aldrich); CaCl_2_ (2 mM, Sigma-Aldrich); NaH_2_PO_4_ (1.2 mM, Sigma-Aldrich); NaHCO_3_ (23 mM, Sigma-Aldrich); and D-Glucose (11 mM, Sigma-Aldrich). The aCSF was adjusted to pH 7.4 using 0.1 mM NaOH (Sigma-Aldrich), was continuously carbogenated, and had an osmolality of ∼300 mOsm. After recovery, slices were left for at least 1 hr at room temperature before recording.

### Electrophysiology, calcium and glutamate imaging

For experiments using calcium and glutamate imaging, mouse hippocampal organotypic brain slices were used. For all other experiments rat hippocampal organotypic brain slices were used. A subset of experiments used acute human cortical brain slices and are specified. Brain slices were transferred to a submerged recording chamber on a patch-clamp rig, which was maintained at a temperature between 28 and 30 °C and were continuously superfused with standard aCSF bubbled with carbogen gas (95 % O_2_: 5 % CO_2,_ Afrox) using peristaltic pumps (Watson-Marlow). The standard aCSF was composed of NaCl (120 mM, Sigma-Aldrich); KCl (3 mM, Sigma-Aldrich); MgCl_2_ (2 mM, Sigma-Aldrich); CaCl_2_ (2 mM, Sigma-Aldrich); NaH_2_PO_4_ (1.2 mM, Sigma-Aldrich); NaHCO_3_ (23 mM, Sigma-Aldrich); and D-Glucose (11 mM, Sigma-Aldrich) in deionized water with pH adjusted to between 7.35 and 7.40 using 0.1 mM NaOH (Sigma-Aldrich). Neurons in the CA3 region of the hippocampus were visualized using a Zeiss Axioskop or Olympus BX51WI upright microscope using 20x or 40x water-immersion objectives and targeted for recording. Micropipettes were prepared (tip resistance between 3 and 7 MΩ) from borosilicate glass capillaries (outer diameter 1.2 mm, inner diameter 0.69 mm, Harvard Apparatus Ltd) using a horizontal puller (Sutter). Recordings were made in current-clamp mode using Axopatch 200B amplifiers (Axon Instruments) and data acquired using WinWCP (University of Strathclyde) or Igor (Markram Laboratory, Ecole polytechnique fédérale de Lausanne). Matlab (MathWorks) was utilised for trace analysis. Basic properties of each cell were recorded (see Figure supplements). Cells were excluded from analyses if the Ra was greater than 80 Ω or if the resting membrane potential was above –40 mV. Two internal solutions were used: a ‘standard’ internal solution (K-gluconate (126 mM), KCl (4 mM), HEPES (10 mM) Na_2_ATP (4 mM), NaGTP (0.3 mM) and Na_2_-phosphocreatine (10 mM); Sigma-Aldrich) and a ‘cesium’ internal solution (CsOH (120 mM), Gluconic acid (120 mM), HEPES (40 mM), Na_2_ATP (2 mM), NaGTP (0.3 mM) and NaCl (10 mM); Sigma- Aldrich). Experimental substances were puffed onto neurons using an OpenSpritzer, a custom-made pressure ejection system (46). Current was injected if required to ensure a neuronal resting membrane potential within 2 mV of -60 mV. In all puffing experiments each data point represents the mean peak puff-induced change in membrane potential from 10 sweeps. For wash-in recordings drugs were washed in for 8 mins before recordings were made. Drugs were washed out for at least 8 mins before wash-out recordings were made. In some experiments tetrodotoxin (TTX) (2 µM, Sigma-Aldrich) was added to the aCSF to block voltage-gated sodium channels.

For calcium and glutamate imaging, organotypic hippocampal mouse brain slices were virally transfected with a genetically-encoded Ca^2+^ reporter (GCAMP6s under the synapsin 1 gene promoter, AAV1.Syn.GCaMP6s.WPRE.SV40, Penn Vector Core) or a genetically-encoded glutamate reporter (iGluSnFR under the human synapsin 1 gene promoter, pAAV.hSynapsin.SF-iGluSnFR.A184S, Addgene) one day post culture using the OpenSpritzer and imaged 5 days later using an Olympus BX51WI microscope, 20x water-immersion objective, CCD camera (CCE-B013-U, Mightex for the calcium imaging and an Andor Zyla 4.2 for the glutamate imaging). For the calcium imaging excitation was provided using a 470 nm LED (Thorlabs). For glutamate imaging, excitation was generated using a LED-based light engine, 475/28 nm (Lumencor). For both imaging paradigms a Chroma 39002 EGFP filter set (480/30 nm excitation filter, 505 nm dichroic, 535/40 nm emission filter) was utilised. Images were collected using µManager (47). Calcium imaging data was analysed using Caltracer3 beta. Glutamate imaging data was analysed using custom scripts written in Matlab.

For real-time imaging of glutamate signaling in neurons adjacent to a live *T. crassiceps* larva, mouse hippocampal organotypic brain slices were virally transfected with a genetically encoded glutamate reporter (iGluSnFR under the human synapsin 1 promoter, pAAV.hSynapsin.SF-iGluSnFR.A184S) one day post culture using the OpenSpritzer. Five to six days later a brain slice would be removed from the culture insert by cutting a roughly 15 mm diameter circle of the membrane out, with the brain slice located in the centre of the circle. The membrane was then placed in a glass-bottomed petri dish containing a drop of aCSF, with the brain slice making contact with the glass bottom. At this point, a single *T. crassiceps* larvae (harvested on the same day or one day prior) of approximately 1 – 2 mm in length was inserted under the membrane with a pasteur pipette and carefully maneuvered until it was adjacent to one edge of the slice. A stainless-steel washer of an appropriate size was then placed on top of the membrane such that it surrounded the brain slice and larva, securing them in place. An additional 2 – 3 ml of carbogen-bubbled aCSF was then added to the petri-dish. The petri-dish was placed in a small incubating chamber (37 °C, 5 % CO_2_) in an LSM 880 airyscan confocal microscope (Carl Zeiss, ZEN SP 2 software). Glutamate signaling in the brain area directly adjacent to the larvae was captured with a tile scan of the area every 90 sec for 15 min using a 40x water-immersion lens (Zeiss) and a 488 nm laser (Zeiss). Transmitted light was also captured to confirm presence and movement of the larva at the site of interest.

Pharmacological manipulations were performed by bath application of drugs using a perfusion system (Watson-Marlow). Mecamylamine, Amiloride, D-AP5 and CNQX were purchased from Tocris. Kynurenic acid and Substance P were acquired from Sigma-Aldrich.

### Data analysis and statistics

Data was graphed and analysed using Matlab, ImageJ, Microsoft Excel and GraphPad Prism. All data was subjected to a Shapiro-Wilk test to determine whether it was normally distributed. Normally distributed populations were subjected to parametric statistical analyses, whilst skewed data was assessed using non-parametric statistical analyses, these included: Mann-Whitney test; Wilcoxon ranked pairs test; Kruskal Wallis one-way ANOVA with post hoc Dunn’s Multiple Comparison test; and Friedman test with post-hoc Dunn’s Multiple Comparison test. The confidence interval for all tests was set at 95%. Raw data source files are available for download from the Open Science page of the Raimondo Lab website: https://raimondolab.com/2024/08/23/data-for-2023-reviewed-preprint-cestode-larvae-excite-host-neuronal-circuits-via-glutamatergic-signaling/

## Acknowledgements

We would like to acknowledge Dr Philip Fortgens (Division of Chemical Pathology, Department of Pathology, University of Cape Town and National Health Laboratory Service (affiliation when the analyses were done) as well as Dr James T. Butler and Dr Roger Melvill of the Epilepsy Unit, Constantiaberg Mediclinic together with two anonymous patients who provided human cortical brain tissue. The research leading to these results has received funding from a Royal Society Newton Advanced Fellowship (NA140170) and a University of Cape Town Start-up Emerging Researcher Award to JVR and grant support from the Blue Brain Project, the National Research Foundation of South Africa, the FLAIR Fellowship Programme (FLR\R1\190829): a partnership between the African Academy of Sciences and the Royal Society funded by the UK Government’s Global Challenges Research Fund, a Wellcome Trust Seed Award and a Wellcome Trust International Intermediate Fellowship (222968/Z/21/Z). KAS was supported by a European Commission Marie Sklodowska-Curie Global Fellowship (Grant 657638, WORMTUMORS). CS, UFP and CPdC were supported by the Federal Ministry of Education and Research of Germany (BMBF), Project title: “CYSTINET-Africa” (01KA1610, Germany II). The funders had no role in study design, data collection and analysis, decision to publish, or preparation of the manuscript.

## Competing interests

The authors declare that there are no competing interests relevant to this manuscript.

## Notes

https://raimondolab.com/2024/08/23/data-for-2023-reviewed-preprint-cestode-larvae-excite-host-neuronal-circuits-via-glutamatergic-signaling/

### Competing Interest Statement

The authors have declared no competing interest.

### Summary of Updates

We have provided multiple revisions in response to reviews provided by eLife, and wish this to be the eLife 'Version of Record'.

## References

1. Diop AG, Boer HM De, Mandlhate C, Prilipko L, Meinardi H. The global campaign against epilepsy in Africa. Acta Trop. 2003;87:149–59.

2. Preux PM, Druet-Cabanac M. Epidemiology and aetiology of epilepsy in sub-Saharan Africa. Lancet Neurol. 2005 Jan;4(1):21–31.

3. Trevisan C, Mkupasi EM, Ngowi HA, Forkman B, Johansen M V. Severe seizures in pigs naturally infected with Taenia solium in Tanzania. Vet Parasitol. 2016;220:67–71.

4. Garcia HH, Del OH. Antiparasitic treatment of neurocysticercosis - The effect of cyst destruction in seizure evolution. Epilepsy & Behavior. 2017;76:158–62.

5. Nash TE, Mahanty S, Garcia HH. Neurocysticercosis-More Than a Neglected Disease. PLoS Negl Trop Dis. 2013;7(4):7–9.

6. Ndimubanzi PC, Carabin H, Budke CM, Nguyen H, Qian YJ, Rainwater E, et al. A systematic review of the frequency of neurocyticercosis with a focus on people with epilepsy. PLoS Negl Trop Dis. 2010;4(11).

7. de Lange A, Mahanty S, Raimondo J v. Model systems for investigating disease processes in neurocysticercosis. Parasitology [Internet]. 2018 Nov 15;1–10. Available from: http://www.ncbi.nlm.nih.gov/pubmed/30430955

8. Garcia HH, Del OH. Antiparasitic treatment of neurocysticercosis - The effect of cyst destruction in seizure evolution. Epilepsy & Behavior. 2017;76:158–62.

9. Vezzani A, Fujinami RS, White HS, Preux P marie, Blümcke I, Sander JW, et al. Infections, inflammation and epilepsy. Acta Neuropathol. 2016;131:211–34.

10. Nash TE, Mahanty S, Loeb J a., Theodore WH, Friedman A, Sander JW, et al. Neurocysticercosis: A natural human model of epileptogenesis. Epilepsia. 2015 Feb;56(2):177–83.

11. Robinson P, Garza A, Weinstock J, Serpa J a, Goodman JC, Eckols KT, et al. Substance P causes seizures in neurocysticercosis. PLoS Pathog [Internet]. 2012 Feb [cited 2013 Sep 11];8(2):e1002489. Available from: http://www.pubmedcentral.nih.gov/articlerender.fcgi?artid=3276565&tool=pmcentrez&rendertype=abstract

12. Sun Y, Chauhan A, Sukumaran P, Sharma J, Singh BB, Mishra BB. Inhibition of store-operated calcium entry in microglia by helminth factors: implications for immune suppression in neurocysticercosis. J Neuroinflammation. 2014 Dec 24;11(1):210.

13. Vendelova E, Camargo de Lima J, Lorenzatto KR, Monteiro KM, Mueller T, Veepaschit J, et al. Proteomic Analysis of Excretory-Secretory Products of Mesocestoides corti Metacestodes Reveals Potential Suppressors of Dendritic Cell Functions. PLoS Negl Trop Dis. 2016;10(10):1–27.

14. Forman CJ, Tomes H, Mbobo B, Burman RJ, Jacobs M, Baden T, et al. Openspritzer: an open hardware pressure ejection system for reliably delivering picolitre volumes. Sci Rep. 2017 May;7(1):2188.

15. Bear MF, Connors BW, Paradiso MA. Neuroscience: Exploring the brain. Third. Lupash E, Connolly E, Dilernia B, Williams P, editors. Philadelphia, Baltimore, New York, London, Buenos Aires, Hong Kong, Sydney, Tokyo: Lippincott Williams & Wilkins; 2007.

16. Bear MF, Connors BW, Paradiso MA. Neuroscience: Exploring the brain. Third. Lupash E, Connolly E, Dilernia B, Williams P, editors. Philadelphia, Baltimore, New York, London, Buenos Aires, Hong Kong, Sydney, Tokyo: Lippincott Williams & Wilkins; 2007.

17. Dingledine R, McBain CJ. Glutamate and Aspartate Are the Major Excitatory Transmitters in the Brain. 1999;

18. Marvin JS, Borghuis BG, Tian L, Cichon J, Harnett MT, Akerboom J, et al. An optimized fluorescent probe for visualizing glutamate neurotransmission. Nat Methods. 2013 Feb;10(2):162– 70.

19. Verastegui MR, Mejia A, Clark T, Gavidia CM, Mamani J, Ccopa F, et al. Novel Rat Model for Neurocysticercosis Using Taenia solium. Am J Pathol. 2015;185(8):2259–68.

20. Stringer JL, Marks LM, White AC, Robinson P. Epileptogenic activity of granulomas associated with murine cysticercosis. Exp Neurol. 2003 Oct;183(2):532–6.

21. Sun DA, Sombati S, DeLorenzo RJ. Glutamate Injury–Induced Epileptogenesis in Hippocampal Neurons. Stroke. 2001 Oct 1;32(10):2344–50.

22. Sombati S, Delorenzo RJ. Recurrent spontaneous seizure activity in hippocampal neuronal networks in culture. J Neurophysiol. 1995 Apr;73(4):1706–11.

23. DeLorenzo R, … SPP of the, 1998 undefined. Prolonged activation of the N-methyl-d- aspartate receptor–Ca2+ transduction pathway causes spontaneous recurrent epileptiform discharges in hippocampal neurons. National Acad Sciences.

24. Anderson WW, Anderson WW, Lewis D V., Scott Swartzwelder H, Wilson WA. Magnesium- free medium activates seizure-like events in the rat hippocampal slice. Brain Res. 1986;398(1):215– 9.

25. Stasheff S, Anderson W, Clark S, Science WW, 1989 undefined. NMDA antagonists differentiate epileptogenesis from seizure expression in an in vitro model. science.sciencemag.org.

26. Ziobro JM, Deshpande LS, DeLorenzo RJ. An organotypic hippocampal slice culture model of excitotoxic injury induced spontaneous recurrent epileptiform discharges. Brain Res. 2011 Jan 31;1371:110–20.

27. Croucher MJ, Bradford HF. NMDA receptor blockade inhibits glutamate-induced kindling of the rat amygdala. Brain Res. 1990;506(2):349–52.

28. Croucher MJ, Bradford HF, Sunter DC, Watkins JC. Inhibition of the development of electrical kindling of the prepyriform cortex by daily focal injections of excitatory amino acid antagonists. Eur J Pharmacol. 1988;152(1–2):29–38.

29. Rice AC, Delorenzo RJ. NMDA receptor activation during status epilepticus is required for the development of epilepsy. Brain Res. 1998;782(1–2):240–7.

30. Hauser WA, Annegers JF, Kurland LT. Prevalence of Epilepsy in Rochester, Minnesota: 1940– 1980. Epilepsia. 1991 Aug;32(4):429–45.

31. Forsgren L, Beghi E, Oun A, Sillanpaa M. The epidemiology of epilepsy in Europe - a systematic review. Eur J Neurol. 2005 Apr 1;12(4):245–53.

32. Ankarcrona M, Dypbukt JM, Bonfoco E, Zhivotovsky B, Orrenius S, Lipton SA, et al. Glutamate-induced neuronal death: A succession of necrosis or apoptosis depending on mitochondrial function. Neuron. 1995 Oct 1;15(4):961–73.

33. Lee HHC, Deeb TZ, Walker JA, Davies PA, Moss SJ. NMDA receptor activity downregulates KCC2 resulting in depolarizing GABAA receptor–mediated currents. Nat Neurosci. 2011 Jun 1;14(6):736–43.

34. Terunuma M, Vargas KJ, Wilkins ME, Ramírez OA, Jaureguiberry-Bravo M, Pangalos MN, et al. Prolonged activation of NMDA receptors promotes dephosphorylation and alters postendocytic sorting of GABAB receptors. Proceedings of the National Academy of Sciences. 2010 Aug 3;107(31):13918–23.

35. Buckingham SC, Campbell SL, Haas BR, Montana V, Robel S, Ogunrinu T, et al. Glutamate release by primary brain tumors induces epileptic activity. Nat Med. 2011 Oct 11;17(10):1269–74.

36. Santosh V, Sravya P. Glioma, glutamate (SLC7A11) and seizures-a commentary. Ann Transl Med. 2017 May;5(10):214.

37. Sontheimer H. A role for glutamate in growth and invasion of primary brain tumors. J Neurochem. 2008 Apr 1;105(2):287–95.

38. Takano T, Lin JHC, Arcuino G, Gao Q, Yang J, Nedergaard M. Glutamate release promotes growth of malignant gliomas. Nat Med. 2001 Sep;7(9):1010–5.

39. Huang W, Choi W, Chen Y, Zhang Q, Deng H, He W, et al. A proposed role for glutamine in cancer cell growth through acid resistance. Cell Res. 2013 May 29;23(5):724–7.

40. Adalid-Peralta L, Arce-Sillas A, Fragoso G, Cárdenas G, Rosetti M, Casanova-Hernández D, et al. Cysticerci drive dendritic cells to promote in vitro and in vivo tregs differentiation. Clin Dev Immunol. 2013;2013:1–9.

41. Verma A, Prasad KN, Cheekatla SS, Nyati KK, Paliwal VK, Gupta RK. Immune response in symptomatic and asymptomatic neurocysticercosis. Med Microbiol Immunol. 2011 Nov 1;200(4):255–61.

42. Prodjinotho UF, Gres V, Henkel F, Lacorcia M, Dandl R, Haslbeck M, et al. Helminthic dehydrogenase drives PGE 2 and IL-10 production in monocytes to potentiate Treg induction . EMBO Rep. 2022 May 4;23(5).

43. Perea G, Araque A. GLIA modulates synaptic transmission. Brain Res Revs. 2010 May;63(1– 2):93–102.

44. Seifert G, Carmignoto G. Astrocyte dysfunction in epilepsy. Brain Res Rev. 2010 May 1;63(1– 2):212–21.

45. Stoppini L, Buchs PA, Muller D. A simple method for organotypic cultures of nervous tissue. J Neurosci Meth. 1991 Apr;37(2):173–182.

46. Forman CJ, Tomes H, Mbobo B, Baden T, Raimondo J V. Openspritzer: an open hardware pressure ejection system for reliably delivering picolitre volumes. bioRxiv. 2016;093633.

47. Edelstein A, Amodaj N, Hoover K, Vale R, Stuurman N. Computer Control of Microscopes Using µManager. In: Current Protocols in Molecular Biology. Hoboken, NJ, USA: John Wiley & Sons, Inc.; 2010. p. 14.20.1–14.20.17.

